# Biological effects of truffle extracts on bacteria, fungi, and microalgae

**DOI:** 10.64898/2025.12.08.692890

**Authors:** N. A. Imidoeva, A. A. Vlasova, E. V. Malygina, A. Yu. Belyshenko, O. E. Lipatova, M. E. Dmitrieva, V. N. Shelkovnikova, E. I. Martynova, T. N. Vavilina, T. Yu. Telnova, M. M. Morgunova, S. S. Shashkina, A. S. Konovalov, V. M. Zhilenkov, D. V. Axenov-Gribanov

**Author notes:** These authors contributed equally to this work (N. A. Imidoeva, A. A. Vlasova).

## Abstract

One of the current challenges on a global scale is the problem of antibiotic resistance. Excessive and inappropriate use of drugs in healthcare and agriculture has accelerated the development of diseases caused by antibiotic-resistant bacteria. In an effort to combat such resistance, it is extremely important to explore alternative sources of new molecules with biological activities. Truffles have become a promising source of new compounds because they contain a wide range of biologically active ingredients. In addition, they are a consortium of symbiotic microorganisms that produce natural products. The aim of this study was to study the antibiotic and growth stimulating effects of methanolic extracts of truffle fruiting bodies collected in Russia in 2020-2022. The extracts demonstrated inhibitory activity against *St. carnosus, E. persicina*, and kanamycin-resistant *E. coli*. Also, for the first time, we have demonstrated that truffle extracts have biological activity in terms of stimulating the growth of *C. vulgaris*. We have observed the effects of short-term and long-term growth stimulation. Thus, truffles can become a promising source of new active ingredients in medicine, agronomy, and different life sciences.

## Introduction

Truffles belong to hypogeous fungi, whose fruiting bodies are formed underground due to mycorrhizal symbiosis with the root systems of plants, primarily oak, pine, hazel, etc. This symbiosis allows truffles to get essential nutrients from host plants. In turn, truffles facilitate absorption of water and minerals from soil by the plant’s root system, thereby promoting the plant growth (Ori *et al*., 2019). One of the primary reasons behind the interest in truffles as a source of new natural products is their ability to interact with a diverse range of microorganisms in their environment. These microbial interactions not only enhance truffle production but also promote synthesis of different natural products and lead to the synthesis of signalling molecules in the truffle fruiting body. Moreover, truffles produce a wide range of secondary metabolites, including sesquiterpenes, triterpenes, polyphenols, and other bioactive compounds with antimicrobial, antioxidant, and anti-inflammatory properties (Dahham *et al*., 2016; Vahdani *et al*., 2017; Wu *et al*., 2022). Also, symbiont bacteria can synthesize volatile organic compounds (VOCs) that contribute to the truffle flavor (Vahdatzadeh *et al*., 2015; Mustafa *et al*., 2020). The VOCs synthesized by truffle symbiont bacteria also play an important role in the interactions with other organisms. For example, a strain of *Staphylococcus aureus* has been shown to synthesize VOCs potentially involved in inhibiting the growth of *T. borchii* mycelium (Barbieri *et al*., 2005). It has been reported that in pure culture, the isolates of *Pseudomonas* spp. obtained from *T. borchii* ascoma are capable of producing phytoregulatory and biocontrol substances that influence the growth and morphogenesis of *T. borchii* mycelium (Sbrana *et al*., 2000).

Nevertheless, the studies of natural product composition in truffles are limited. Given that truffles and their symbionts represent an understudied group, they have the potential to produce novel natural products. For example, research indicates that certain components found in truffles can inhibit growth of different pathogenic and non-pathogenic microorganisms. In the study by Casarica *et al*. (2016), it was shown that aqueous extract of desert truffles of *Terfezia claveryi* species had antimicrobial activity against *E. coli* ATCC 8739, *St. aureus* ATCC 6538P and *S. epidermidis* ATCC 12228. The study performed by Tejedor-Calvo *et al*. (2021) reveals the antimicrobial properties of extracts of the desert truffle *Tirmania nivea*. In the mentioned study, phenolic compounds are capable of inhibiting the growth of seven types of described bacteria. However, it is noteworthy that the antimicrobial activity of Russian black and white truffles has not been studied before.

In addition to the association between truffles and bacteria, cyanobacteria and microalgae are also regular members of the truffle consortium (Menotta *et al*., 2004). One of the well-studied groups of microalgae belongs to genus *Chlorella. Chlorella* sp. is a microalgae that is widely used as a source of pharmacologically active metabolites with different biological activities (Fedorov *et al*., 2013; Hamouda *et al*., 2022). This microalga is also being studied for couplings with other organisms, namely bacteria and viruses. For example, it has been shown that *Azospirillus* sp. and *Bacillus* sp. can stimulate growth of *C. vulgaris*. In the study by Cho K. *et al*. (2019), it was shown that co-cultivation of algae and bacteria stimulates the growth of *C. vulgaris* and increases its biomass.

Thus, the aim of our study was to investigate the antibiotic and growth stimulating properties of black and white truffles of the genus *Tuber*.

## Materials and methods

### Truffle samples

Fruiting bodies of truffles were collected in Krasnodar region (Russia) between August and November in 2020‒2022. Thus, in 2020, truffle fruiting bodies were collected in Krasnodar (black truffles: N = 2). In 2021, samples were taken from Krasnodar (black truffles: N =5, white truffles: N = 2) and Sochi (black truffles: N =3). In 2022, samples of fungi analyzed were obtained from Krasnodar (black truffles: N =12) and Maykop (black truffles: N =4). All delivery procedures were carried out in plastic containers filled with soil and rice to prevent rotting during transportation. The truffles were transported under thermostatic conditions at 6‒10 °C. The received samples were cleaned from soil with a brush, washed and dried, and stored in the fridge at ‒20 °C until further experiments. The summary on the state of truffles and place of their sampling is presented in Table 1.

**Table 1.**
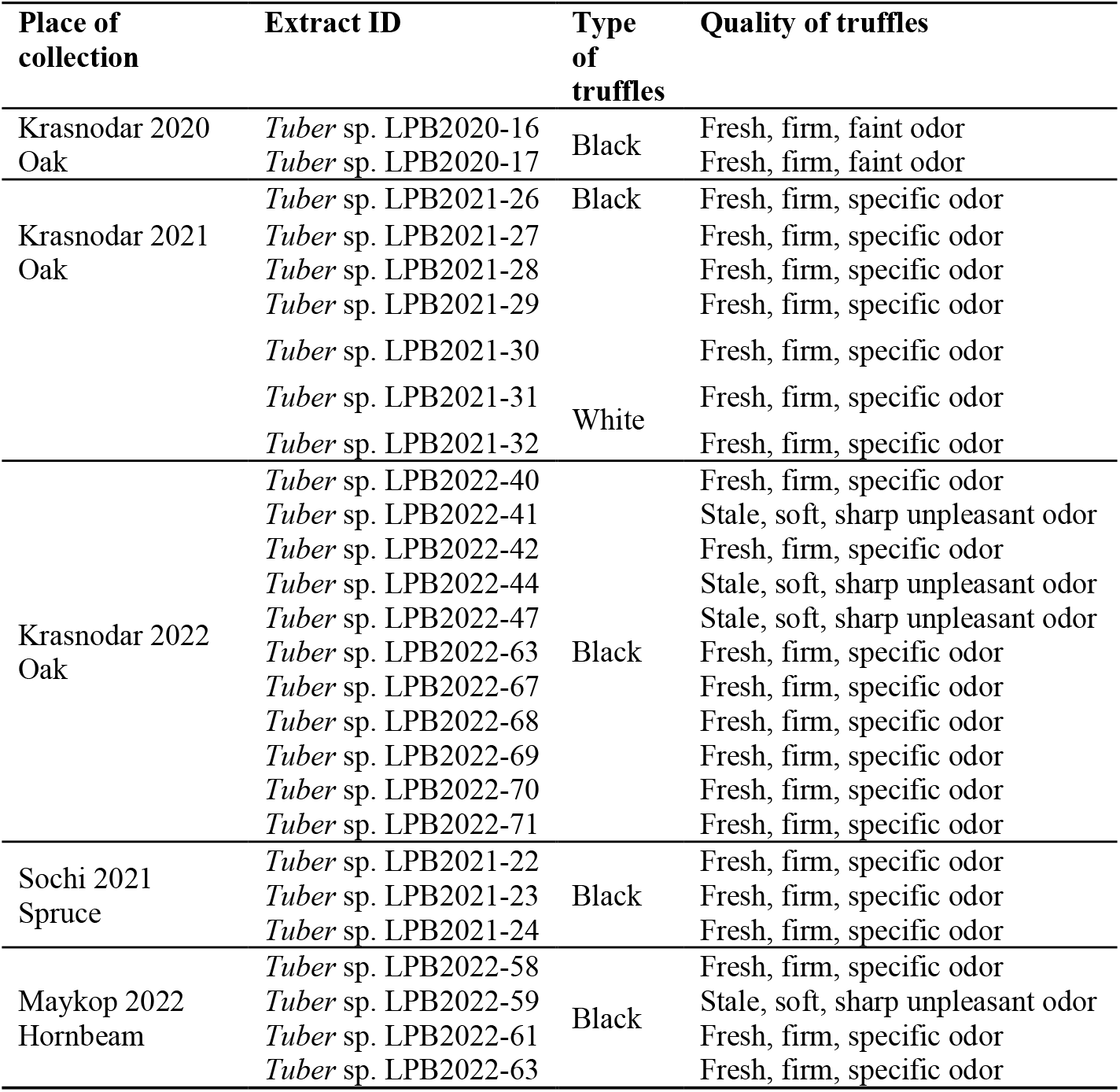
List of extracts, obtained from mushrooms and tested in this study.

Wild black truffles were sampled in several cities of the Southern region of the Russian Federation between August and November in 2020‒2022. The samples were taken near the cities of Krasnodar, Maykop, and Sochi.

Fungi sized 3 to 5 cm were collected and cleaned with a soft toothbrush and then sterilized with 70% ethyl alcohol. Then, the truffles were dried and shock-frozen in liquid nitrogen. Until further experiments, the samples were stored in liquid nitrogen

### Extraction

To perform the extraction, 0.5 g of each truffle sample were defrosted and homogenized using a mortar and pestle with methanol added in a ratio of 1:10. The mixture was incubated and shaken for one hour on a roller shaker MX-T6-S (China). Then, the samples were centrifuged at 3 000 rpm for 10 min using DM 0412 centrifuge (Hettich, USA). The extracts were dried in-vacuo and re-dissolved in the concentration of 10 mg/mL (Angelini *et al*., 2014). Each truffle extract was assigned an identification number in Table 1.

### Antibiotic activity of truffle extracts

The antibiotic activity was tested using the disk diffusion method (Ruangpan, 2004). Several strains of microorganisms were chosen as model test cultures. The 24-hour test cultures were inoculated on different solid nutrient media and dried at room temperature for up to 40 min. To assess the antibiotic activity, 45 µL of extracts were loaded onto 5 mm paper disks and dried at room temperature. Then the disks were placed onto Petri dishes with inoculated and dried test cultures. The plates with disks were incubated for 24 hours. Inhibition zones were measured with ±1 mm accuracy. Disks containing only methanol, served as negative controls (Axenov-Gribanov *et al*., 2016).

In this study, we used 18 test strains representing a wide range of gram-positive and gram-negative bacteria, as well as fungi, which allowed comprehensive assessment of the antibacterial and antifungal properties of truffles. Table 2 presents the list of test cultures, media and temperature of cultivation.

**Table 2.** Test strains and conditions of cultivation.

| Test cultures | Nutrient media | Temperature of cultivation |
| --- | --- | --- |

| Gram-positive |  |  |
| --- | --- | --- |
| *Staphylococcus carnosus* B-8726 | LB | 37 °C |
| *Bacillus subtilis* ATCC 66337 | LB | 37 °C |
| *Bacillus subtilis* B-4537 | MPA | 30 °C |
| *Bacillus subtilis* 168 | LB | 37 °C |
| *Micrococcus luteus* B-7846 | YEPD | 30 °C |
| *Mycobacterium smegmatis* AC-1339 | ISP | 37 °C |
| *Staphylococcus aureus* B-6646 | MPA | 37 °C |
| *Kocuria rhizophila* B-5389 | MPA | 30 °C |
| *Enterococcus mundtii* B-12673 | LB | 37 °C |
| *Staphylococcus carnosus* B-8726 | LB | 37 °C |
| Gram- negative |  |  |
| *Pseudomonas putida* B-4589 | LB | 30 °C |
| *Erwinia persicina* 19328 | LB | 30 °C |
| *Pseudomonas aeruginosa* B-6643 | MPA | 37 °C |
| *Escherichia coli* B-6645 | MPA | 37 °C |
| *Escherichia coli* lptD CamR | LB | 37 °C |
| *Escherichia coli* tolC KanR | LB | 37 °C |
| Yeast |  |  |
| *Saccharomyces cerevisiae* Y-1173 | YPD | 30 °C |
| *Candida utilis* Y-3089 | YEPD | 30 °C |
| *Candida albicans* Y-3108 | YEPD | 37 °C |

The composition of media was as follows: LB media (tryptone—10 g/L, yeast extract—5 g/L, NaCl—5 g/L); MPA media (nutritious dry broth based on enzymatic beef meat hydrolysate – 30 g/L, peptone – 9 g/L); YEPD media (glucose – 20 g/L, yeast extract – 5 g/L, peptone – 10 g/L); ISP media (peptone – 5 g/L, yeast extract – 3 g/L, malt extract – 3 g/L, glucose – 10 g/L); YPD media (soy peptone – 20 g/L, yeast extract – 10 g/L, sucrose – 20 g/L).

#### Biological activity of truffle extracts on algae

In this study, the biological activity of extracts was evaluated on the basis of growth of *C. vulgaris*. First, the mother culture of algae was grown in 50% of Tamiya nutrient medium at 36 °C for 5 days (Ördög *et al*., 2012; Kalinina *et al*., 2023). Then, the culture was diluted in 50% of Tamiya nutrient media until the optical density value reached 0.06-0.08 at the wavelength of 560 nm. General technique of spectrophotometric analysis was applied using PE-5300V spectrophotometer (PromEcoLab, Russia). Composition of 100% Tamiya nutrient medium is as follows: KNO_3_ - 5.0 g/L, MgSO_4_ х 7H2O - 2.5 g/L, KH_2_PO_4_ х 3H_2_O - 1.25 g/L, FeSO_4_ х 7H_2_O - 0.009 g/L, H_3_BO_3_ - 2.86 g/L, MnCl_2_ х 4H_2_O - 1.81 g/L, ZnSO_4_ х 7H_2_O - 0.22 g/L, MoO_3_ х 4H_2_O – 0.018 g/L, NH_4_VO_3_ - 0.023 g/L (Grigoriev, 2014).

For the experiment, aliquots of methanol extracts in the amount of 100 µl were added to conical flasks. The solvent was evaporated, and then, 40 mL of the *C. vulgaris* culture was added to the dry residue. The control group of *C. vulgaris* was cultivated with dry residue of methanol added, as a solvent of truffle extract. Truffle extracts were not added to the samples.

The experiment was conducted in three stages. In the first stage, the experimental group included the following extracts: *Tuber* sp. LPB2020-16, *Tuber* sp. LPB2020-17, *Tuber* sp. LPB2021-22, *Tuber* sp. LPB2021-23, *Tuber* sp. LPB2021-24, *Tuber* sp. LPB2021-26, *Tuber* sp. LPB2021-28, *Tuber* sp. LPB2021-29, and *Tuber* sp. LPB2021-27. In the second stage, the experimental group included the following extracts: *Tuber* sp. LPB2022-58, *Tuber* sp. LPB2022-59, *Tuber* sp. LPB2022-61, *Tuber* sp. LPB2022-67, and *Tuber* sp. LPB2022-69. In the third stage, the experimental group included the following extracts: *Tuber* sp. LPB2021-30, *Tuber* sp. LPB2022-39, *Tuber* sp. LPB2022-42, and *Tuber* sp. LPB2022-47. Each stage had its own control group of *C. vulgaris*.

Further, control and experimental groups were cultivated at 36 °C using whirlpool photothermoshaker x12-1 (Mycotech, Russia). The cultivation time was up to 6 days with 900 Lux light flow. To assess the dynamics of the algae growth, we measured the culture optical density on days 2, 4 and 6 of the experiment (Hamouda *et al*., 2022).

### Statistics

In total, we estimated the effects of 18 fruiting bodies of truffles. All experiments were carried out in 3–7 biological replicates. Statistical analysis was performed with Past software (V. 4.03) using Tukey’s Proc ANOVA test. Tukey’s range test, p-value of ≤0.05 was considered to indicate statistical significance (Maurício *et al*., 2023).

## Results

### Antibiotic activity of truffle extracts

Methanolic extracts from truffles showed antibiotic activity against only three out of 18 test strains studied: *St. carnosus* B-8726, *E. persicina* 19328, and *E. coli* tolC KanR (Table 3).

**Table 3.**
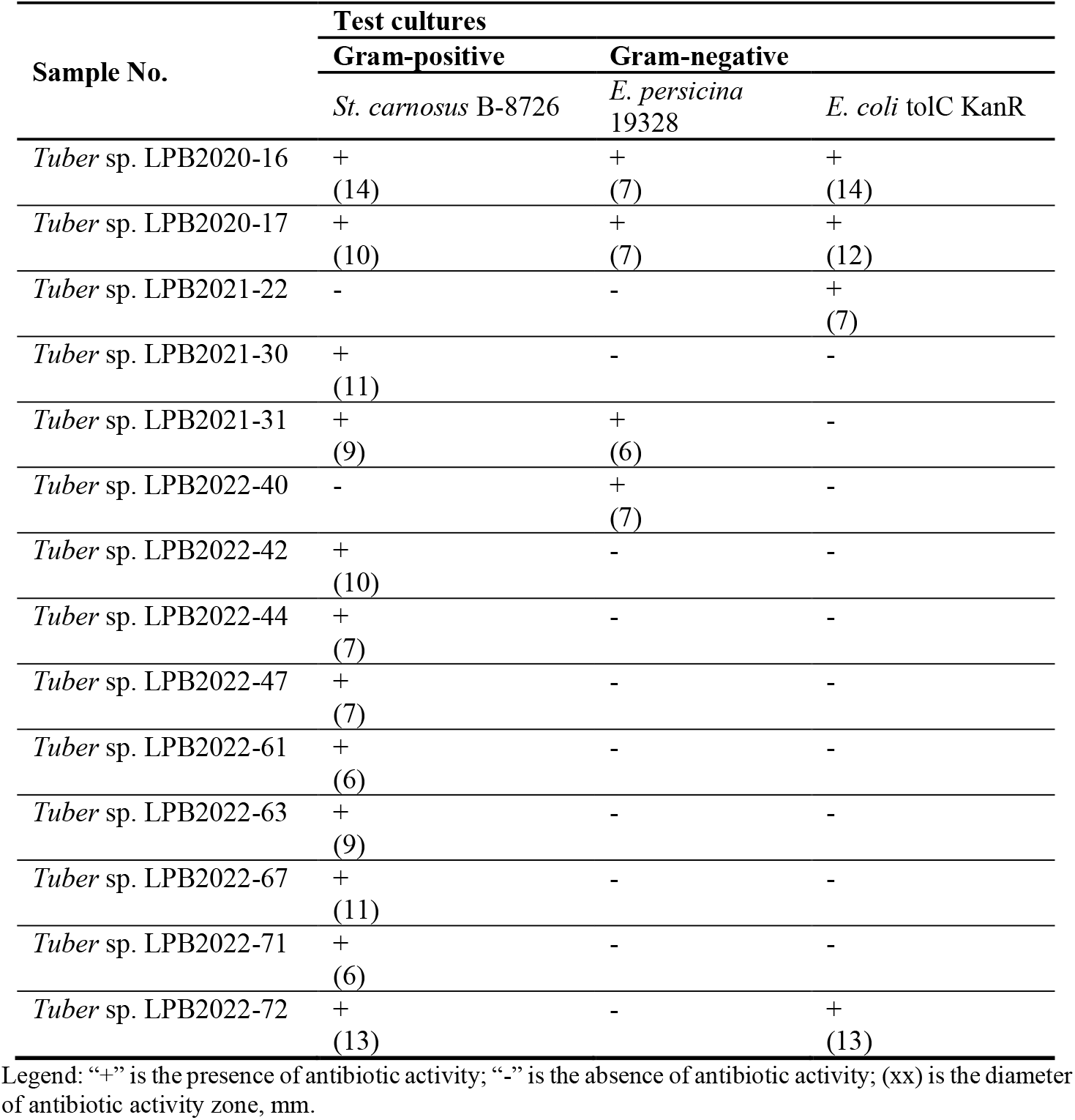
Antibiotic activities of fruiting bodies of truffle methanolic extracts.

| Sample No. | Test cultures |  |  |
| --- | --- | --- | --- |
|  | Gram-positive | Gram-negative |  |
|  | *St. carnosus* B-8726 | *E. persicina* 19328 | *E. coli* tolC KanR |
| *Tuber* sp. LPB2020-16 | +(14) | +(7) | +(14) |
| *Tuber* sp. LPB2020-17 | +(10) | +(7) | +(12) |
| *Tuber* sp. LPB2021-22 | - | - | +(7) |
| *Tuber* sp. LPB2021-30 | +(11) | - | - |
| *Tuber* sp. LPB2021-31 | +(9) | +(6) | - |
| *Tuber* sp. LPB2022-40 | - | +(7) | - |
| *Tuber* sp. LPB2022-42 | +(10) | - | - |
| *Tuber* sp. LPB2022-44 | +(7) | - | - |
| *Tuber* sp. LPB2022-47 | +(7) | - | - |
| *Tuber* sp. LPB2022-61 | +(6) | - | - |
| *Tuber* sp. LPB2022-63 | +(9) | - | - |
| *Tuber* sp. LPB2022-67 | +(11) | - | - |
| *Tuber* sp. LPB2022-71 | +(6) | - | - |
| *Tuber* sp. LPB2022-72 | +(13) | - | +(13) |

Legend: “+” is the presence of antibiotic activity; “-” is the absence of antibiotic activity; (xx) is the diameter of antibiotic activity zone, mm.

Thirteen out of 28 fruiting bodies, including 1 white and 12 black truffles, inhibited growth of gram-positive bacteria *St. carnosus*. Another interesting finding was that only four out of 28 fruiting bodies showed activity against *E. persicina*. Furthermore, it was demonstrated that truffle methanolic extracts showed antibiotic activity against kanamycin-resistant *E. coli*, while exhibiting no activity against typical *E. coli*.

### Biological activity of truffle extracts on algae

During the experiment, growth of *C. vulgaris* was estimated on days 2, 4 and 6 of the experiment. Figure 1 demonstrates the growth of *C. vulgaris* when methanolic extracts of truffles were added. Also, Table 4 shows the dynamics of growth of *C. vulgaris* algae (expressed in % related to control group).

**Table 4.** Daily growth dynamics of the microalga *C. vulgaris*.

| Sample No. | 2-day | 4-day | 6-day |
| --- | --- | --- | --- |
| *Tuber* sp. LPB2020-16 | 2 | 3 | 7 |
| *Tuber* sp. LPB2020-17 | 8 | 9 (*) | 9 (*) |
| *Tuber* sp. LPB2021-26 | 28 (*) | 18 (*) | 13 (*) |
| *Tuber* sp. LPB2021-27 | 24 (*) | 7 (*) | -3 |
| *Tuber* sp. LPB2021-28 | 26 (*) | 13 (*) | 8 |
| *Tuber* sp. LPB2021-29 | 12 | 4 | 0 |
| *Tuber* sp. LPB2021-30 | 15 (*) | 10 (*) | 2 |
| *Tuber* sp. LPB2022-67 | 15 (*) | 7 (*) | 7 (*) |
| *Tuber* sp. LPB2022-69 | 15 (*) | 7 (*) | 2 |
| *Tuber* sp. LPB2022-39 | 20 (*) | 15 (*) | 4 (*) |
| *Tuber* sp. LPB2022-42 | 6 (*) | 3 | 2 |
| *Tuber* sp. LPB2022-47 | 5 (*) | 8 (*) | 6 (*) |
| *Tuber* sp. LPB2021-22 | 6 | 9 (*) | 8 (*) |
| *Tuber* sp. LPB2021-23 | 39 (*) | 14 (*) | 8 |
| *Tuber* sp. LPB2021-24 | 16 (*) | 19 (*) | 13 |
| *Tuber* sp. LPB2022-58 | 4 | 0 | -3 |
| *Tuber* sp. LPB2022-59 | 9 (*) | 0 | 0 |
| *Tuber* sp. LPB2022-61 | 15 (*) | 8 (*) | 3 |

Legend: Asterisks indicate significant values. Red color means long-term stimulation effects; Yellow color — short-term effects. Black color — no effects

**Figure 1.**
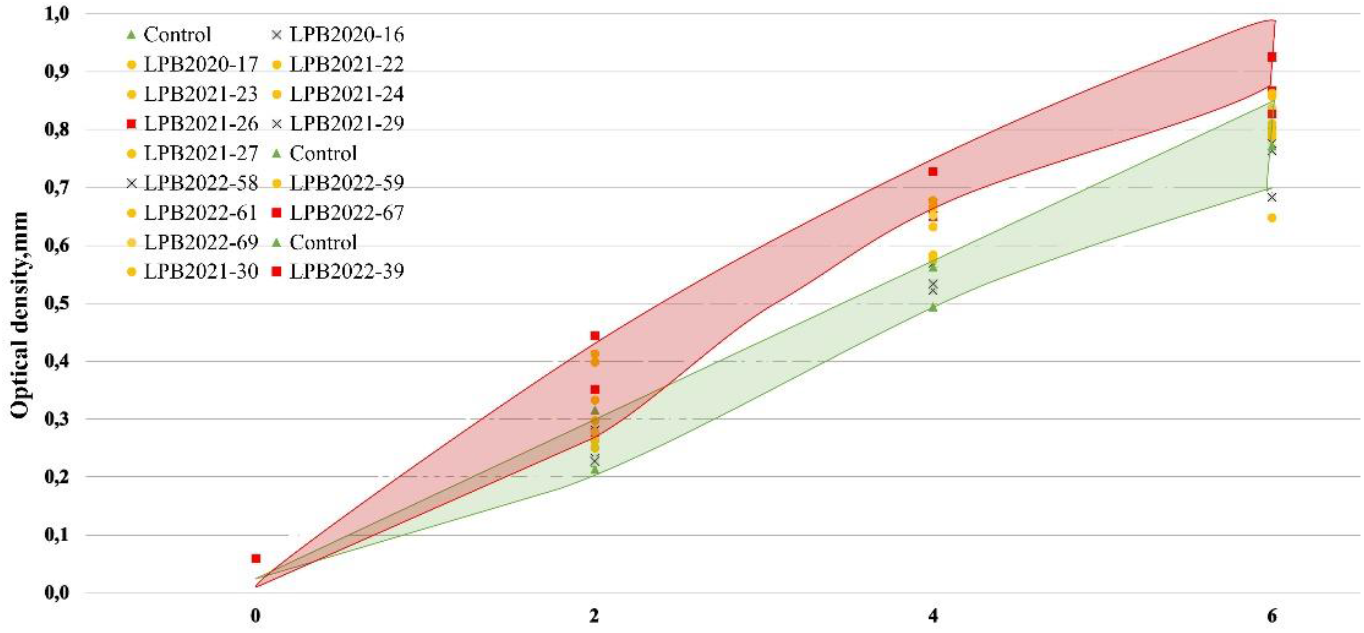
Optical density of *C. vulgaris* in control and experimental conditions. Legend: ▲ - Control; ● - significant differences observed in one or two control points (short-term effects); █ - significant differences observed during 6 days of experiment (long-term effects); ✖ - absence of significant differences. P-value ⩽0.05

Green area in Figure 1 indicates the area of the *C. vulgaris* control group cultivated without adding truffle extracts. Red area shows the zone where the experimental group showed notable differences from the control one, throughout the experiment (long-term effects). Yellow markers show the short-term stimulation effects on the algae, but significant differences from the control group can be seen in one or two control points. Gray markers indicate extracts that did not stimulate growth of *C. vulgaris*.

Four out of 18 truffle extracts exhibited the highest long-term stimulating effect on algal growth. During 6 day of the experiment, we observed significant stimulating effects of the following methanol extracts: *Tuber* sp. LPB2021-26, *Tuber* sp. LPB2022-67, *Tuber* sp. LPB2022-39, and *Tuber* sp. LPB2022-47. After adding *Tuber* sp. LPB2021-26 extract to the culture of *C. vulgaris*, we observed the increase of the algal culture optical density. Optical density of the algal suspension increased from 0.06 to 0.352 on the 2nd day of the experiment, 0.494 on the 4th day, and 0.868 on the 6th day. Compared to control conditions, in the experimental group, optical density of the algal culture was 56%, 36% and 26% higher on the 2nd, 4th and 6th days of the experiment, relatively. In the case of the effect of *Tuber* sp. LPB2022-67 extract, the daily growth reached 30% in the first two days of experiment and then decreased to the end of experiment (Table 4).

Also, it was found that some extracts exerted partial stimulation (short-term stimulation effects) of algal growth. Significant differences between control and experimental groups were observed at one or two points in time. For example, truffle extract *Tuber* sp. LPB2021-22 demonstrated significant differences on days 4 and 6 of the experiment. Growth of *C. vulgaris* comprised 7.5% to 8.5% per day. Extracts of *Tuber* sp. LPB2021-23 and *Tuber* sp. LPB2021-24 stimulated the algae growth and demonstrated significant differences compared to control conditions during the first four days of the experiment. On the 2nd day, the optical density of *Tuber* sp. LPB2021-24 extract was 0.298, which was 32% higher than under control conditions. On day 4, optical density was 0.678 (37% rate of stimulation).

The *Tuber* sp. LPB2021-23 extract revealed the highest short-term stimulation activity towards the algae. Optical density of *Tuber* sp. LPB2021-23 extract at the 2nd day point was 0.399, which was 77% higher related to the control conditions. However, by day 6 of the experiment, the effects of the truffle extract decreased, and optical density of algal culture was similar to the control value.

The study revealed that adding extracts of *Tuber* sp. LPB2020-16, *Tuber* sp. LPB2021-29, and *Tuber* sp. LPB2022-58 did not stimulate algal growth.

## Discussion

Thus, it has been demonstrated that methanolic extracts of truffle are characterized by inhibitory effects against bacteria, and growth-stimulating effects against microalgae. Three out of 18 bacterial and fungal cultures were inhibited by the extracts. These strains were represented by *St. carnosus, E. persicina*, and *E. coli* resistant to the antibiotic kanamycin.

*St. carnosus* is a common strain found on almost any surface, including soil, plants, and animals (Sommerfeld *et al*., 2023). Some strains of *St. carnosus* are known to cause human diseases, such as osteomyelitis, infective endocarditis, infectious arthritis (Mohammad *et al*., 2022). Antibiotic activity against this strain can also be of value in pharmaceutical industry to develop new antimicrobial agents and drugs.

*E. persicina* is a pathogenic bacterium that primarily affects agricultural plants by causing leaf spot and necrosis on its host (Canik Orel *et al*., 2023). Antibiotic activity of truffle’s methanolic extract against the pathogen *E. persicina* may prove beneficial in addressing the issue of controlling the spread of this pathogen and protecting crops from its effects.

The study revealed that methanolic extracts of truffles displayed antibiotic activity specifically against kanamycin-resistant strain *E. coli*. Resistance to kanamycin is mediated by the tolC gene, which encodes a protein involved in the efflux of the antibiotic from the cell. This strain of bacteria is used in research, it allows assessment of the effectiveness of antibiotics (Guo *et al*., 2013). Since kanamycin-resistant *E. coli* has a mutation in the tolC gene, it is more susceptible to antimicrobial properties of the truffle’s extract compared to typical test-strain of *E. coli*. The mutation in the tolC gene may have weakened the bacteria’s ability to expel the antibiotic compounds from the truffle extract, leading to inhibition of their growth. This specificity of antibiotic activity highlights the potential of truffles as a novel antimicrobial agent against bacterial strains resistant to traditional antibiotics.

In addition, the research findings indicate that methanolic extracts of fruiting bodies of truffles, collected in 2021 and 2022, exhibit low antibiotic activity compared to those collected in 2020. Comparisons were made by measuring the zones of inhibition of the extracts against microorganisms or test cultures. This means that the truffles collected in 2020 have not lost their properties after 3 years of storage at ‒20 °C. Perhaps this is due to the fact that the truffles were collected from the same locations in the Krasnodar region. A possible reason for this is that over time, intensive truffle harvesting can deteriorate the soil composition, leading to a decline in concentration of important minerals and nutrients. Low soil quality can have adverse effects on truffle development, and result in reduced antimicrobial properties. In other cases, there was no connection between the year and the place where the fruiting bodies were collected.

Also, the antibiotic activity of extracts of truffle fruiting bodies against only three out of 18 test strains can be attributed to several factors, including specific pathogen ‒ host interactions, variability in extract composition and species-specific susceptibility. Antibiotic activity observed in the majority of test strains may stem from their inherent ability to withstand the effects of truffle extracts. Understanding of these factors is crucial to harness the full antimicrobial potential of truffle extracts and to explore the potential of future applications in agronomy, medical fields, and food safety.

Also, materials of these study reveal that extracts from truffles can stimulate the growth of *C. vulgaris*, demonstrating the potential beneficial applications of truffles beyond their antimicrobial properties and found no direct correlation between the stimulating effects on algae and the state of truffles, place of their sampling, or their quality. At the same time, several reactions of *C. vulgaris* were observed when cultivating it with truffle extracts. We demonstrated that the highest stimulating effects of extracts occurred during the first two days of the experiment. Subsequently, all types of truffle extracts retained their stimulating effect on *C. vulgaris*, but a decrease in growth rate was noted in Table 4. This could be explained by composition and energetic value of the nutrient media that decreased during the experiment. Another explanation could be that over time *C. vulgaris* caused a thinning of the culture medium and reduced the amount of stimulating natural products in the truffle extracts. In this case, the variable effects of our extracts may be related to unstable chemical composition of the truffles due to unstable content of microflora associated with them.

As mentioned above, activity of the extracts can be related to the microflora inhabiting truffles. The life cycle of truffles includes symbiosis with not only host plants, but also with microorganisms. Symbiont bacteria exert their influence not only on truffles themselves, but also on the surrounding organisms, too (Menta and Pinto, 2016). Therefore, we hypothesize that it was specific symbiont bacteria that had the resulting effect on the growth of the alga *C. vulgaris*.

Volatile organic compounds produced by microorganisms can influence plant adaptability, they can also interfere with plant-animal interactions. By doing so, these compounds promote plant growth and prevent the emergence of harmful microbes (Barbieri *et al*., 2005; Junker and Tholl, 2013). This may suggest that the volatile organic compounds produced by truffle symbiont bacteria may stimulate growth of *C. vulgaris*. The microbial composition of Russian black truffles collected in Sochi has been described earlier in our studies (Malygina *et al*., 2024). The primary analysis detected at least 45 bacterial organisms with OTUs, and 22 fungal organisms as regular inhabitants of *Tuber* sp. fruiting bodies (Monaco *et al*., 2020; Morgunova *et al*., 2023).

Different composition of symbionts of truffles, influencing the content of natural products in the metabolome of truffles, allows us to hypothesize that both truffles and microalgae could have common natural products in their metabolomic pathways. These natural products can be found in literature, they are 1-octen-3-ol (eight-carbon-containing volatiles) and 2-methyl-1-propanol. As predicted before, these natural products have been detected in most truffles species and might act as signals to plants (Li *et al*., 2023). Also, 1-octen-3-ol previously was found in *Chlorella* sp., as a product of linoleic acid oxidation and was described as a natural product responsible for aroma of seafood (Selli and Cayhan, 2009; Zhang *et al*., 2009; Nader *et al*., 2022).

The findings from this study underscore the dual role of truffles as both antimicrobial agents and growth stimulators in different biological systems. The observed antibiotic activities against specific bacteria, especially those resistant to conventional treatments, suggest that truffles could be a valuable resource in the search for new antimicrobial agents. At the same time, the growth promotion seen in *C. vulgaris* highlights the potential applications of truffles in agricultural and ecological contexts. Future research should focus on elucidating the mechanisms underlying these effects, including the roles of specific bacterial symbionts, to better harness the full potential of truffles in both medical and agricultural fields. Continued exploration of truffle metabolites and their interactions with microorganisms may provide deeper insights into how these fungi can contribute to sustainable practices in crop protection and the development of new therapeutic strategies.

In summary, the study on the biological effects of methanolic extracts from truffles has revealed significant insights into their antibiotic and growth-stimulating properties. We examined the antibiotic activity of methanolic extracts of 28 fruiting bodies of black and white truffles against 18 different test-cultures of bacteria and fungi. The results showed that the extracts were effective against *St. carnosus, E. persicina*, and *E. coli* resistant to the antibiotic kanamycin. The analysis also revealed that the truffles collected in 2021 and 2022 probably had low antibiotic activity compared to those collected in 2020 in Krasnodar, possibly due to soil degradation from intensive harvesting. Also, the study found no direct correlation between the antimicrobial and growth-stimulating effects and the quality or sampling location of the truffles. This suggests that the bioactive compounds may be influenced by a complex interplay of environmental and biological factors. Overall, the findings of this study position truffles as a promising resource for developing new pharmaceutical substances and agricultural applications, paving the way for future research aimed at harnessing their full potential in combating antibiotic resistance and enhancing crop protection.

## Acknowledgement

This work was supported by the [Russian Science Foundation] under Grant [22-76-10036].

